# Ventral hippocampal CA1 and CA3 differentially mediate learned approach-avoidance conflict processing

**DOI:** 10.1101/180513

**Authors:** Anett Schumacher, Franz R. Villaruel, Sadia Riaz, Andy C. H. Lee, Rutsuko Ito

## Abstract

The ventral hippocampus is thought to play a key role in the resolution of approach-avoidance conflict, a scenario that arises when stimuli with opposing valences are present simultaneously. Little is known, however, about the contributions of specific hippocampal sub-regions in this process, a critical issue given the functional and anatomical heterogeneity of this structure. Using a non-spatial cue-based paradigm in rats, we found that transient pharmacological inactivation of ventral CA1 produced an avoidance of a conflict cue imbued with both learned positive and negative outcomes, whereas inactivation of the ventral CA3 resulted in the opposite pattern of behavior, with significant preference for the conflict cue. In contrast, dorsal CA1- and CA3- inactivated rats showed no change in conflict behavior. Our findings provide important insight into the functions and circuitry of the ventral hippocampus by demonstrating that the ventral CA1 and CA3 subserve distinct and opposing roles in approach-avoidance conflict processing.

**Grant sponsor:** Natural Science and Engineering Research Council (N.S.E.R.C) of Canada awarded to RI (402642) and ACHL (402651).

## INTRODUCTION

The regulation and successful resolution of approach-avoidance conflict is a ubiquitous dilemma that organisms commonly face. Deciding to approach or avoid requires evaluating the incentive value of environmental stimuli that may be associated with both positive and negative valences. These ambivalent stimuli evoke a state of motivational conflict, which needs to be resolved in order that an effective response can be executed to maintain survival (Miller, 1944).

Despite the prevailing view that the predominant function of the hippocampus (HPC) is in mnemonic processing and/or spatial cognition (O'Keefe and Nadel, 1978; Eichenbaum, 2000), a significant body of rodent work has pointed towards a role for this structure in processing approach-avoidance conflict (for recent review see Ito and Lee, 2016). Animal models of approach-avoidance conflict have typically involved the initial establishment of an approach response, followed by the induction of conflict by later punishing the same learned response (Geller and Seifter, 1960; Vogel et al., 1971) or alternatively, taking advantage of conflicting innate behaviours (e.g. desire to explore vs. fear of being in an exposed environment) in ethological tests of anxiety such as the open field test and the elevated plus maze (e.g. Lister, 1990; Rodgers et al., 1997; Belzung and Griebel, 2001). HPC lesions typically cause persistence of learned approach responses in the face of punishment (e.g. Kimura, 1958; Isaacson and Wickelgren, 1962; Kimble, 1963), and an increase in approach behaviour in ethological tests of anxiety (e.g. Bannerman et al., 1999; McHugh et al., 2004; Trivedi and Coover, 2004). This is also consistent with a substantial body of work showing potentiation of behavioural indices of appetitive motivation in HPC-lesioned animals, which include feeding, cued approach, intracranial self-stimulation and progressive schedules of food reinforcement (Davidson and Jarrard, 1993; Tracy et al., 2001; Ito et al., 2005; Davidson et al., 2009; 2013). Together, these data speak to the HPC, and particularly the ventral aspect of this structure having a critical role in the suppression of approach responses in situations of uncertainty (Gray and McNaughton, 2000; McHugh et al., 2008; Abela et al., 2013; Bannerman et al., 2014; Schumacher et al., 2016), and a threat to energy balance (Tracy et al., 2001). Convergent with this, a growing number of neuropsychological and functional neuroimaging studies in humans have demonstrated that the human analogue of the rodent ventral HPC, the anterior HPC, is significantly involved when participants are required to respond or make decisions under scenarios of high conflict (Bach et al., 2014; O'Neil et al., 2015; Oehrn et al., 2015; Loh et al., 2017).

Crucially, while much work on the role of the HPC in approach-avoidance conflict has focused on its differentiation along the dorsal-ventral axis of this structure, there has been comparatively little insight into potential functional differences along the transverse axis. The dentate gyrus (DG), CA3 and CA1 are three distinct subregions along this axis, and form predominantly unidirectional excitatory circuits, from DG to CA3, and then to CA1 (Amaral and Witter, 1989; Van Strien et al., 2009). While these subregions can be clearly demarcated on the basis of anatomical, physiological and computational evidence, separating them on the basis of functional evidence has been less straightforward. Existing research exploring the functional dissociations of HPC subfields have largely focused on their role in memory encoding and retrieval, novelty detection and spatiotemporal processes within the dorsal HPC, and attribute overlapping functions to the subregions. The DG has been implicated in novelty/mismatch detection, memory encoding and the ability to discriminate two similar memory representations (i.e. pattern separation). On the other hand, CA1 and CA3 have been implicated in varying degrees of memory encoding, pattern separation and completion (e.g. retrieval of a complete mnemonic representation based on a partial or degraded input) depending on the nature of information and/or degree of dissimilarity between mnemonic representations and incoming sensory information (Lee et al., 2004; Leutgeb et al., 2004; e.g. Vazdarjanova and Guzowski, 2004; Hoge and Kesner, 2007; Barbosa et al., 2012). Notably, substantially fewer studies have specifically explored the differential functions of these subregions within the ventral HPC. Those that have, have implicated ventral CA3 (vCA3) in spatial and non-spatial novelty detection, and retrieval of contextual fear memory, whereas ventral CA1 (vCA1) has been demonstrated to be crucial for the temporal ordering of olfactory information, non-spatial novelty detection and retrieval of contextual fear (Hunsaker et al., 2008; Beer et al., 2014). Moreover, of particular relevance to the present work, recent research has highlighted a critical role for ventral DG (vDG) in ethological tests of anxiety, with lesions to this region resulting in increased time spent in the open arms of the elevated plus maze and the central region of the open field maze in comparison to similar manipulations to dorsal DG (Weeden et al., 2015). To our knowledge, however, the involvement of vCA1 and vCA3 in approach-avoidance behaviour has yet to be explored systematically.

The current study sought, therefore, to reveal the differential contributions of rodent vCA1 and vCA3, and dorsal CA1 (dCA1) and CA3 (dCA3) to approach-avoidance conflict processing. To achieve this, we used a novel cue-based approach-avoidance paradigm (Figure 1) that has been recently shown to be sensitive to ventral HPC damage (Schumacher et al., 2016). In contrast to traditional rodent models of approach-avoidance conflict, a key strength of this task is that it is non-spatial in nature, an important characteristic given the role of the HPC in spatial cognition. Moreover, our paradigm is unique in that it uses learned, as opposed to innate, appetitive and aversive cues, and is able to disentangle the acquisition of incentive values from the expression of motivational conflict. Rats first learned to associate three distinct tactile cues with a positive, negative or neutral outcome. Post-acquisition GABA_A_ and GABA_B_ agonist microinfusions using a muscimol/baclofen (M/B) cocktail were then conducted to selectively inactivate the CA1 or CA3 regions of the dorsal or ventral HPC, prior to an approach-avoidance conflict test in which the appetitive and aversive cues were presented in combination to create motivational conflict, alongside the neutral cue. We report that vCA1 inactivation induced a potentiation of avoidance tendency from the conflict cue, whereas vCA3 inactivation induced the opposite pattern, of increased approach tendency to the conflict cue. In contrast, dCA 1 or dCA3 inactivation had no impact on conflict behavior. Our results implicate ventral HPC subregions in having bidirectional control over approach-avoidance behaviour in the face of motivational conflict.

**Figure 1.**
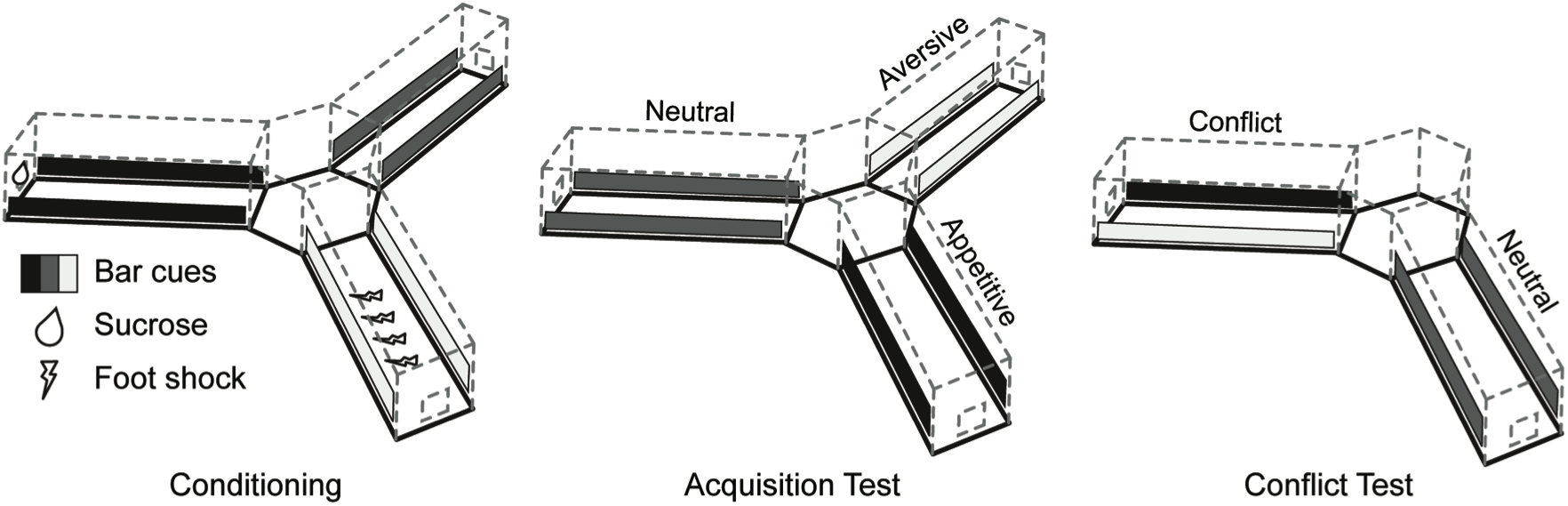
Schematic diagram of the non-spatial cue-based learned approach-avoidance paradigm. There were four different phases: **Habituation** (not shown), in which rodents were exposed to the radial arm apparatus and visuospatial cues. **Conditioning**, in which rodents learned the outcomes (appetitive, aversive, or neutral) associated with three visotactile cues. To minimise the use of spatial information, the positions of the cues were changed across conditioning sessions, with the maze rotated left or right by varying degrees (60°, 120°, or 180°) between each session, and the entire maze was covered with red cellophane film to block the visibility of extra-maze cues. **Acquisition test**, in which rodents were assessed on their learning of the outcomes associated with each cue. **Conflict Test**, in which rodents were presented with a superimposition of positive and negative cues in one arm, and a neutral cue in another arm.

## RESULTS

### Histological Verifications

Inactivation sites of the dCA1 and dCA3 ranged from -3.3 to -3.8mm posterior to bregma (Paxinos and Watson, 1998), whereas rats with ventral HPC infusions showed inactivation sites ranging from -4.8 to -6.04mm posterior to bregma (Figure 2). Five rats in the dCA1 and dCA3 groups and 8 rats in the vCA1 and vCA3 groups had infusion sites outside of the targeted regions and were, therefore, excluded from the study. Furthermore, a total of 6 rats did not acquire the mixed valence conditioning, and were subsequently removed from the study. Final group numbers were as follows: The ventral HPC group contained a total of 38 rats with 18 receiving M/B injections: 9 vCA1(M/B), 9 vCA3(M/B); and 20 receiving saline only injections as a control comparison: 10 vCA1(SAL) and 10 vCA3(SAL). The dorsal HPC group consisted of 23 rats with 6 dCA1(M/B), 6 dCA3(M/B), 5 dCA1(SAL), and 6 dCA3(SAL). It is worth noting that there was neither extensive damage to the neuronal tissue nor extended gliosis around the injection site, indicating accurate surgical technique and infusion into the desired HPC subregion.

**Figure 2.**
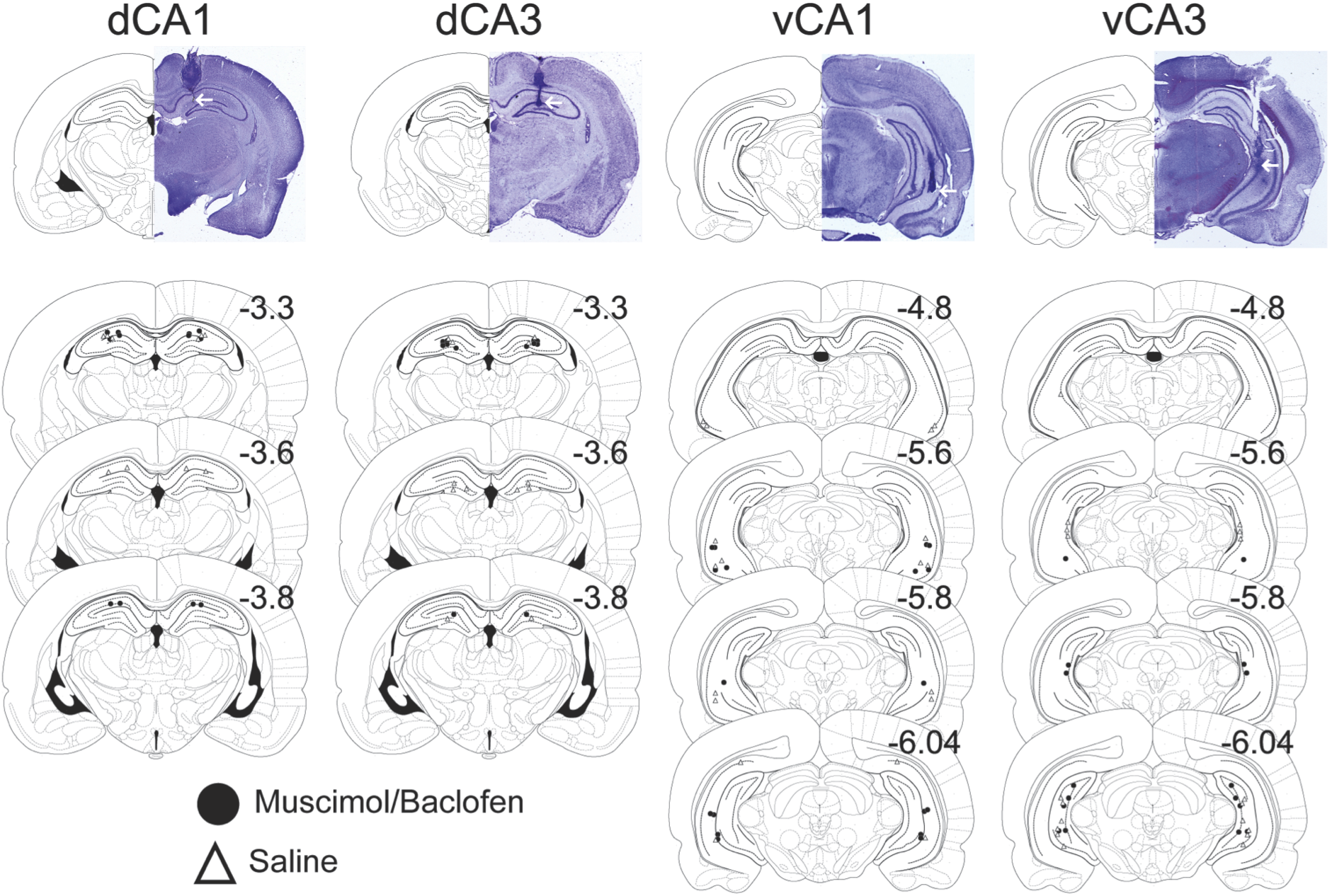
Representative photographic and schematic diagrams showing the location of the injector tips relative to bregma in the dorsal and ventral CA1 and CA3 of the hippocampus, for rats included in the data analysis. Black circles represent animals that received muscimol/baclofen at test and white triangles represent rats that received saline at test.

### Approach-Avoidance Conflict

#### Habituation

Rats underwent three habituation sessions, one of which mimicked the conditions of the final approach-avoidance conflict test. More specifically, rats were exposed to two maze arms, one containing the neutral cue and another containing a superimposition of cues that were eventually assigned as appetitive and aversive cues. There were no differences in the time spent exploring the two arms (cues) during this habituation session in any of the dorsal HPC groups (Arm: F(1, 33) = 0.75, p = 0.40) or ventral HPC groups (Arm: F(1,33) = 0.36, p = 0.55). Nor were there any significant differences in the exploratory performance of rats assigned to the dorsal HPC groups (Arm x Drug x Group: F(1,19) = 0.27, p = 0.61) or ventral HPC groups (Arm x Drug x Group, F(1, 33) = 1.73, p = 0.20).

#### Acquisition tests

Rats performed a total of nine conditioning sessions to associate non-spatial cues with an appetitive, aversive or neutral outcome. Learning was assessed by performing a conditioned cue approach-avoidance test after 4 and 8 conditioning sessions, *without* any drug/saline infusions. ANOVA of the time spent in each of the three arms in the two cue acquisition tests revealed that all rats in the dorsal HPC group acquired the cue-outcome associations successfully by Test 2 (Arm: F(2, 38) = 43.22, p < 0.0001; Test: F(1, 19) = 7.77, p = 0.012; Arm x Test interaction: F(2, 38) = 11.31, p < 0.0001), with rats spending significantly more time in the appetitive arm (dCA1(M/B): p < 0.02; dCA1(SAL): p < 0.04; dCA3(M/B): p < 0.001; dCA3(SAL): p < 0.05) and less time in the aversive arm (dCA1(M/B): p < 0.01; dCA1(SAL): p < 0.01; dCA3(M/B): p < 0.01; dCA3(SAL): p < 0.04), relative to the neutral arm (Figure 3A). In addition, there were no pre-existing group differences in the acquisition of the three cue-outcome relationships (Drug: F(1, 19) = 0.49, p = 0.49; Region: F(1, 19) = 0.015, p = 0.90; Test x Arm x Drug x Region: F(2, 38) = 0.331, p = 0.72).

Similarly, ANOVA of the time spent in each of the three arms in the two acquisition tests in the four ventral HPC groups revealed that all rats acquired the cue-outcome associations successfully by Test 2 (Arm: F(2, 66) = 102.58, p < 0.0001; Arm x Test interaction: F(2, 66) = 5.19, p < 0.01), as evidenced by their spending more time in the appetitive arm (vCA1(M/B): p < 0.01; vCA1(SAL): p < 0.001; vCA3(M/B): p < 0.03: vCA3(SAL): p < 0.01) and less time in the aversive arm (vCA1(M/B): p < 0.001; vCA1(SAL): p < 0.02; vCA3(M/B): p < 0.001; vCA3(SAL): p < 0.001), relative to the neutral arm (Figure 3B). In addition, there were no preexisting group differences in the acquisition of the three cue-outcome associations (Region: F(1, 33) = 1.19, p = 0.28; Drug: F(1,33) = 1.23, p = 0.28; Test x Arm x Drug x Region: F(2,66) = 0.58, p = 0.53).

**Figure 3.**
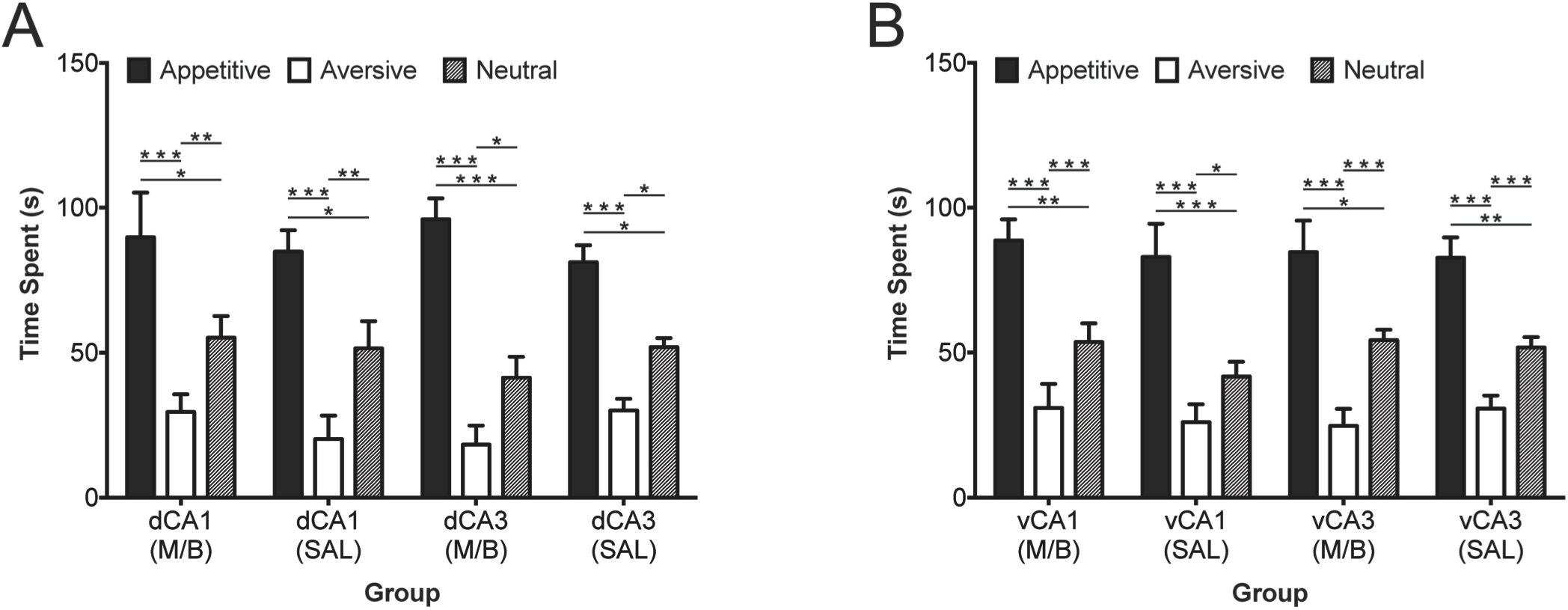
Mean (± SEM) time spent in arms with “appetitive,” “aversive,” and “neutral” cues during a test of concurrent conditioned cue preference and avoidance for rats that received either muscimol/baclofen (M/B) or saline (SAL) at test in the (A) dorsal CA1 and CA3 hippocampus, and the (B) ventral CA1 and CA3 hippocampus. * p < 0.05, ** p < 0.01, *** p < 0.001. Mean and standard error values, as well as 95% confidence intervals are reported in Supplemental Table S1.

### Conflict Test

#### Conflict cue approach-avoidance

The conflict test was administered following the successful acquisition of the three cue-outcome associations. The test session involved the rats being allowed to freely explore two arms: combined appetitive and aversive cues in one arm (conflict arm), and neutral cues in another arm (neutral arm), following bilateral microinfusions of drug M/B or saline into target sites. ANOVA of the overall time spent in each of the two arms revealed that rats in all four dorsal HPC groups showed no difference in the time spent exploring the two arms during the conflict test (Arm: F(1,19) = 0.01, p = 0.94, h_p_^2^ = 0.0001), indicating that neither approach, nor avoidance of the conflict cue dominated their behaviour in the face of motivational conflict (Figure 4A). Furthermore, the performance of the dCA1- and dCA3-inactivated groups did not significantly differ from that of the saline controls (Arm x Drug x Region, F(1, 19) = 1.73, p = 0.21, h_p_^2^ = 0.08).

In contrast, ANOVA of the overall time spent in each of the two arms in the four ventral HPC groups revealed significantly altered performance in the conflict test between the vCA1 and vCA3-inactivated groups, and their control groups (Arm x Drug x Region interaction: F(1, 33) = 15.30, p < 0.0001, h_p_^2^ = 0.32, Figure 4B). More specifically, simple main effects analyses revealed a significant main effect of Arm in the vCA1-inactivated (F(1, 33) = 12.71, p < 0.001, h_p_^2^ = 0.28) and vCA3-inactivated (F(1, 33) = 21.68, p < 0.0001, h_p_^2^ = 0.40) groups, but not in either of the saline groups (vCA1: F(1,33) = 0.0001, p = 1.0, h_p_^2^ = 0.0001, vCA3: F(1, 33) = 0.64, p = 0.43, h_p_^2^ = 0.019), indicating that the vCA1-inactivated rats spent significantly less time in the conflict arm (p < 0.001) while vCA3-inactivated rats spent more time in the conflict arm (p < 0.001), compared to that in the neutral arm. Furthermore, inactivated vCA1 rats spent significantly more time in the neutral arm compared to their saline control group (p < 0.001), while vCA3-inactivated rats showed a decreased time spent in the neutral arm compared to their saline control group (p = 0.034). Thus, vCA1 inactivation led to increased avoidance tendencies, while vCA3 inactivation led to increased approach tendencies in the face of motivational conflict.

**Figure 4.**
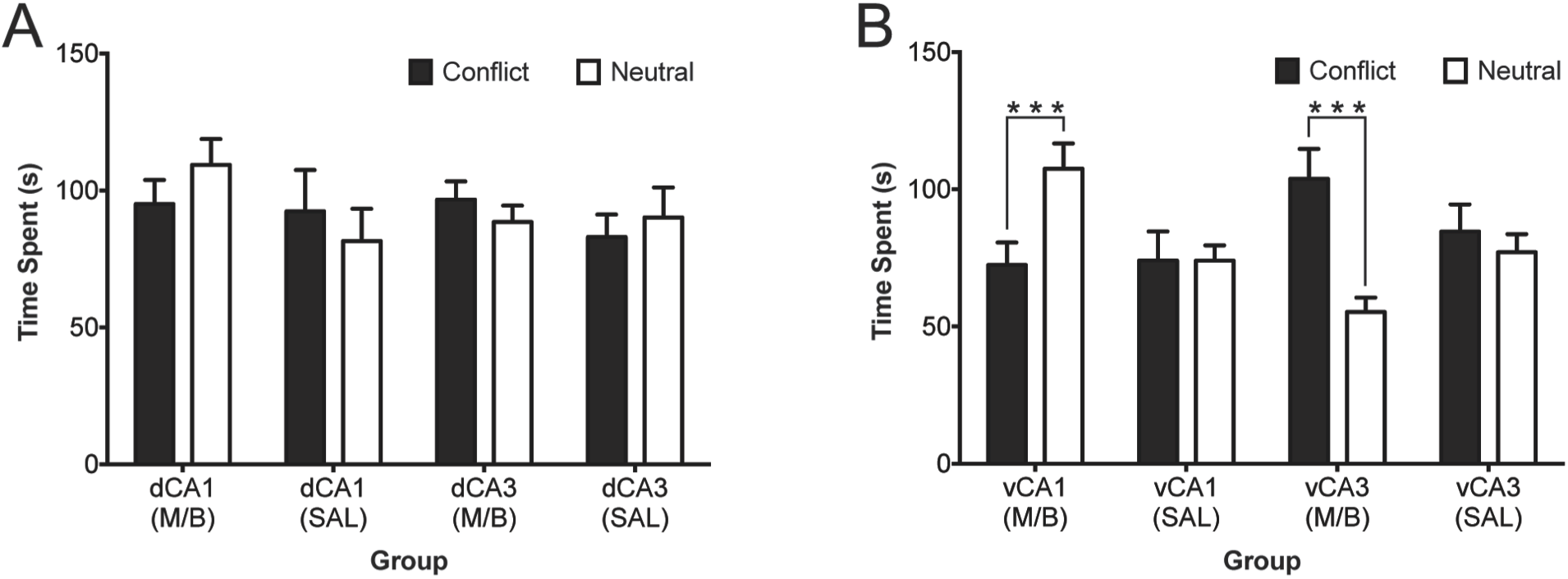
Mean (± SEM) time spent in arms containing cues of conflicting valence (superimposed appetitive and aversive cues) or neutral cues during the conflict test following administration of muscimol/baclofen (M/B) or saline (SAL) in the (A) dorsal CA1 and CA3 hippocampus, and in the (B) ventral CA1 and CA3 hippocampus. Ventral CA1 inactivation lead to significantly more time spent in the neutral than the conflict arm, while ventral CA3 inactivation lead to significantly more time spent in the conflict arm than the neutral arm. *** p < 0.001. Mean and standard error values, as well as 95% confidence intervals are reported in Supplemental Table S2.

#### Number of entries into conflict and neutral arms

ANOVA of the total number of full body entries made into the conflict and neutral arms during the conflict test in all dorsal HPC groups (Fig 5A) revealed no significant effects of any kind (Arm (F(1, 19) = 1.86, p = 0.95; Group: F(1, 19) = 1.54, p = 0.36; Arm x Region x Drug F(1, 19) = 0.016, p = 0.9). In contrast, ANOVA of the total number of full body entries made into the conflict and neutral arms during the conflict test in all ventral HPC groups (Fig 5B) revealed a significant Arm x Region x Drug interaction (F(1, 33) = 5.05, p = 0.031) as well as a significant Arm x Drug interaction (F(1,33) = 7.09, p = 0.012). Subsequent simple effects analyses and pairwise comparison analyses attributed the significant three-way interaction to the number of entries between the conflict and neutral arms being significantly different in vCA1(p = 0.03) and vCA3 drug groups (p = 0.02), with the vCA1-inactivated rats making fewer entries into the conflict arm, and the vCA3-inactivated rats making more entries into the conflict arm, compared to the neutral arm.

**Figure 5.**
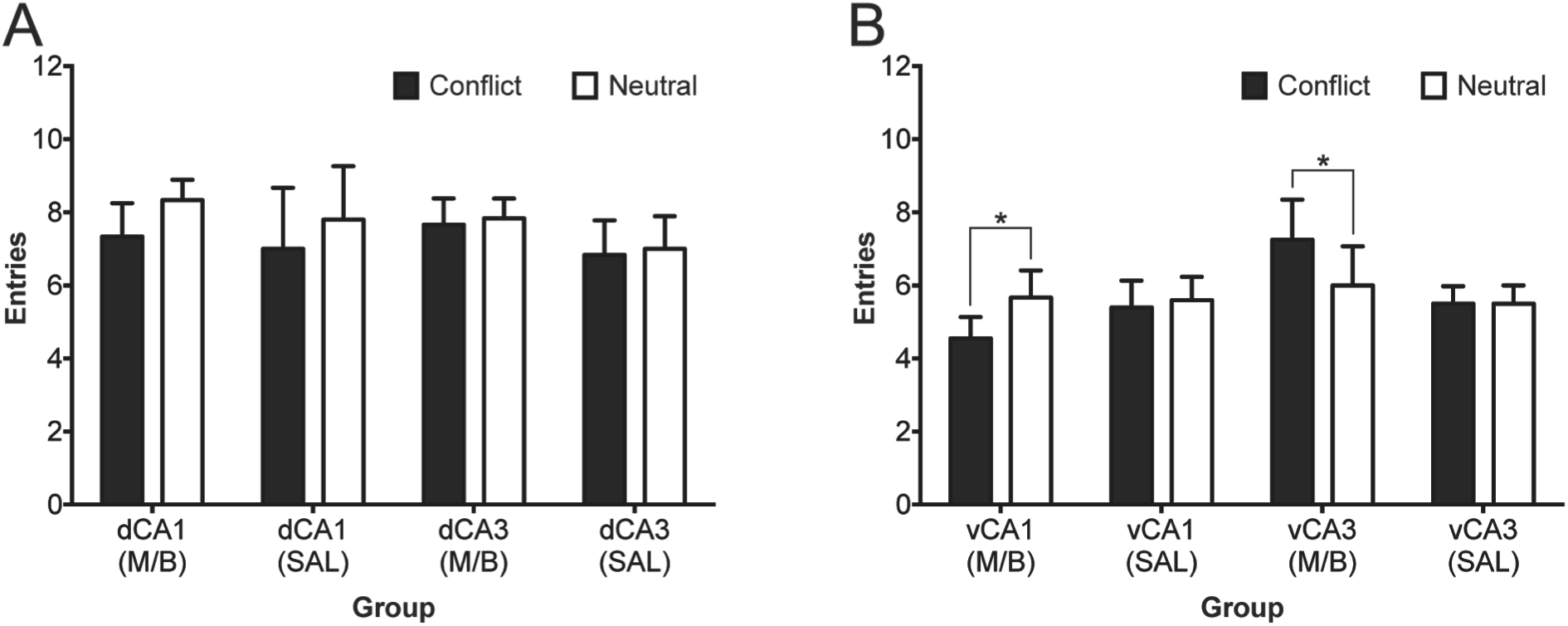
Mean (± SEM) number of entries in arms containing cues of conflicting valence (superimposed appetitive and aversive cues) or neutral cues during the conflict test following administration of muscimol/baclofen (M/B) or saline (SAL) in the (A) dorsal CA1 and CA3 hippocampus, and in the (B) ventral CA1 and CA3 hippocampus. Mean and standard error values, as well as 95% confidence intervals are reported in Supplemental Table S2.

### Light Dark Box

A standard ethological test of anxiety, the light dark box task, was used to assess potential differences in anxiety levels. ANOVA of the time spent exploring the light vs. dark compartments of the box in the dorsal HPC group (Figure 6A) revealed that rats spent significantly more time in the dark, compared to the light compartment (Compartment: F(1(19) = 34.32, p < 0.001), and that there was no difference in performance between groups (Drug x Region: F(1, 19) = 2.26, p = 0.15; Compartment x Drug x Region, F(1, 19) = 0.11, p = 0.75). ANOVA of the time spent in the dark vs. light compartments in the ventral HPC groups (Figure 6B) revealed a significant Drug x Compartment interaction (F(1, 33) = 9.69, p < 0.01), but no other significant main effects, nor interactions (all p > 0.05). Further simple effects analyses revealed the significant interaction effect to be attributable to the ventral HPC-inactivated groups (vCA1(M/B) and vCA3(M/B)) collectively spending significantly more time in the light compartment, and less time in the dark compartment than their saline counterparts (p < 0.01). Thus, ventral HPC inactivation only, irrespective of the subfield targeted, induced a reduction in anxiety.

**Figure 6.**
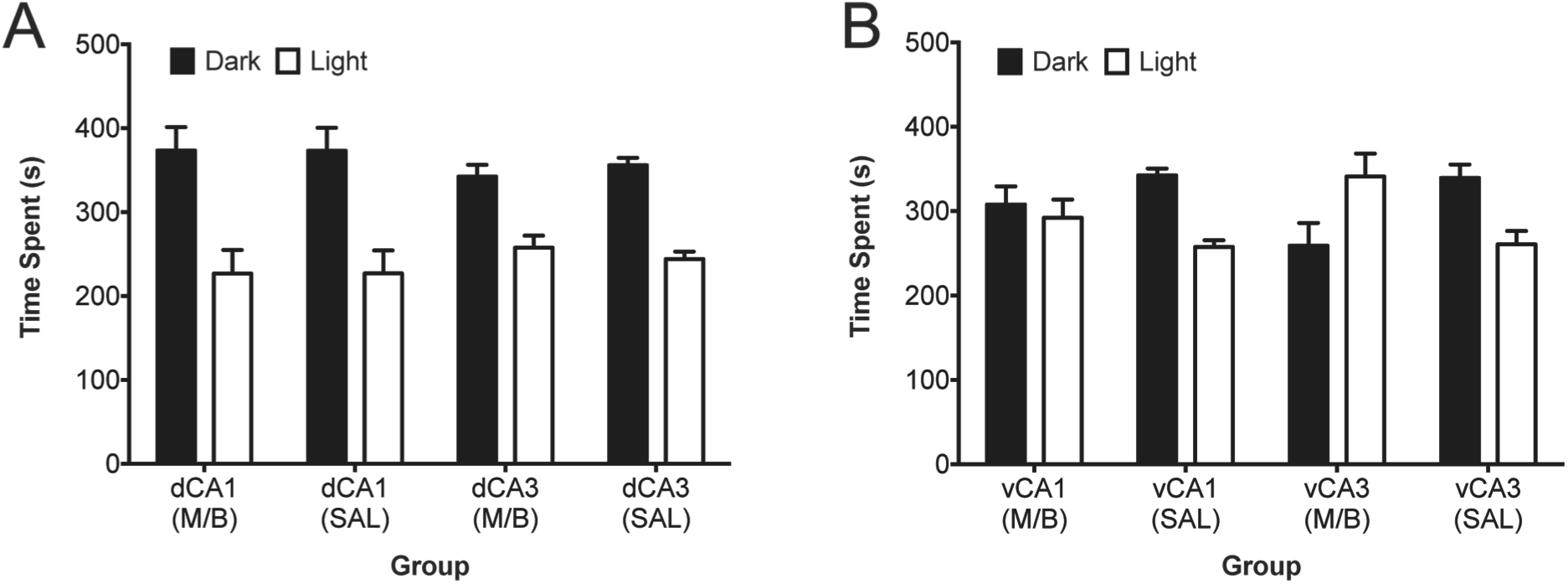
Mean (± SEM) time spent in the dark and light compartments during a light-dark box test of anxiety following administration of muscimol/baclofen (M/B) or saline (SAL) in the (A) dorsal CA1 and CA3 hippocampus, and in the (B) ventral CA1 and CA3 hippocampus. Mean and standard error values, as well as 95% confidence intervals are reported in Supplemental Table S3.

### Novelty Detection Test

To rule out alternative explanations of the conflict test data, rats were administered a novelty detection test in the same radial arm maze as that used for the approach-avoidance conflict paradigm, in which they were first allowed to explore two ‘familiar’ arms and were then exposed to a third ‘novel’ arm. ANOVA of the time spent exploring the novel vs. familiar arms in the dorsal HPC-manipulated groups (Figure 7A) revealed a significant Arm x Drug interaction (F(1, 19) = 14.13, p < 0.0001) and a main effect of Arm (F(1, 19) = 25.07, p < 0.0001), but no threeway Region x Drug x Arm interaction (F(1,19) = 0.07, p = 0.80). Simple effects analysis exploring the significant Arm x Drug interaction revealed that the saline-infused groups (dCA1(SAL) and dCA3 (SAL)) spent significantly more time in the novel arm, relative to the familiar arm (< 0.0001). In contrast, rats with inactivated dCA1 and dCA3 spent equal time exploring the familiar and novel arms (p = 0.38). Furthermore, rats inactivated in the dCA1 and dCA3 spent more time in the familiar arm compared to their respective control groups (p < 0.0001). Similarly, ANOVA of the time spent exploring the novel vs. familiar arms in the ventral HPC-infused groups (Figure 7B) revealed a significant Arm x Drug interaction (F(1, 30) = 18.6, p < 0.0001) and main effect of Arm (F(1,30) = 42.22, p <0.0001), but no three-way Region x Drug x Arm interaction (F(1, 30) = 0.21, p = 0.65). Subsequent simple effects analysis revealed the significant Arm x Drug interaction to be due to the saline-infused groups (vCA1(SAL) and vCA3 (SAL)) spending significantly more time in the novel arm, relative to the familiar arm (p < 0.0001), whereas there was no difference in time spent between the familiar and novel arms for both the vCA1(M/B) and vCA3(M/B) groups (p = 0.14). Furthermore, inactivation of the vCA1 and vCA3 led to less time spent in the novel arm, relative to their control groups (p < 0.0001). Thus, inactivation of CA1 and CA3 regions in both the ventral HPC and dorsal HPC resulted in impaired exploration of the novel arm.

**Figure 7.**
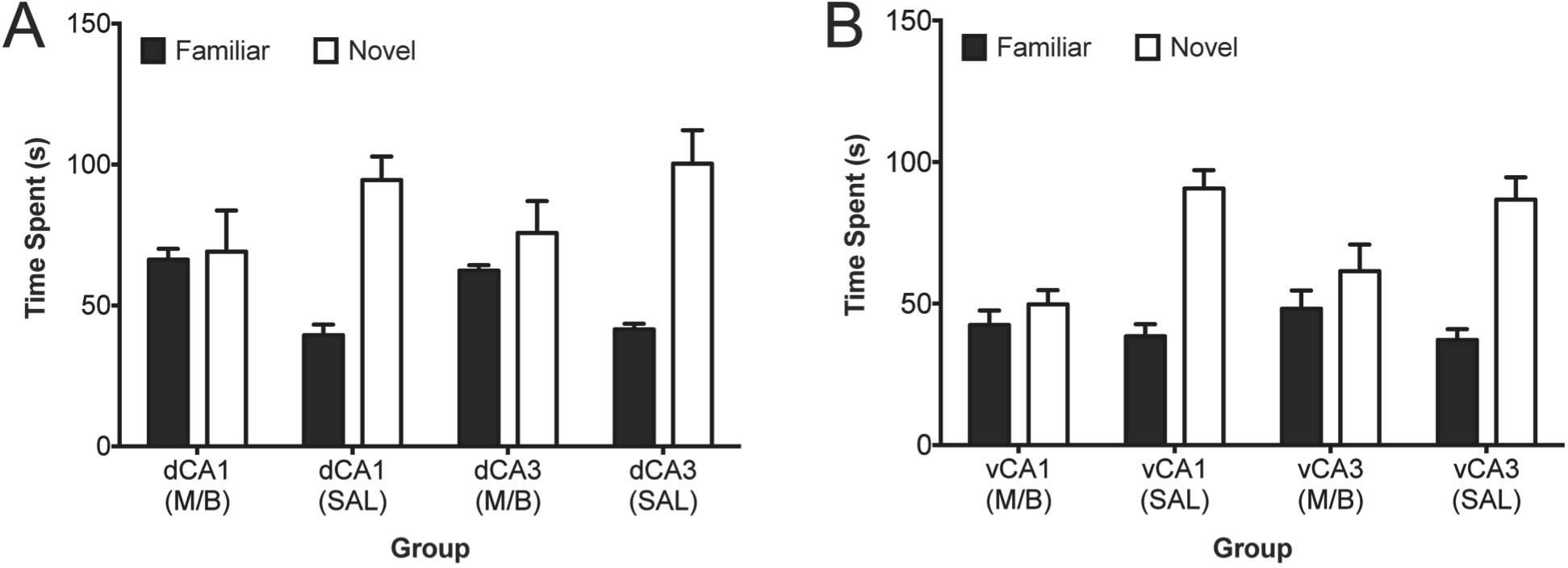
Mean (± SEM) time spent in a novel and familiar arm during a test of spatial novelty following administration of muscimol/baclofen (M/B) or saline (SAL) in the (A) dorsal CA1 and CA3 hippocampus, and the (B) ventral CA1 and CA3 hippocampus. Mean and standard error values, as well as 95% confidence intervals are reported in Supplemental Table S4.

### Locomotor Activity

Finally, baseline locomotor activity (Figure 8) was also measured to ensure that potential differences in exploration times in the approach-avoidance conflict tests were not confounded by changes in general activity. As expected, there was a significant within-subjects effect of Bin (F(11,209) = 10.86, p < 0.0001) reflecting the fact that there was a decrease of locomotor activity across all groups. However, there was no significant difference in spontaneous locomotor activity between the dorsal HPC groups (Region: F(1,19) = 1.71, p = 0.27, Region x Drug x Bin: F(11,209) = 0.60, p = 0.82). The ventral HPC rats yielded similar results: there was no overall Region effect (F(1,31) = 0.11, p = 0.75) or any significant 3- or 2- way interactions between Region, Bin and Drug (all F > 0.07, p > 0.12) but there was a significant within-subjects effect of Bin (F(11, 341) = 10.38, p < 0.0001) reflecting decreased locomotor activity over time. In summary, no differences in the baseline activity were found between animals of different subregions of the dorsal HPC and ventral HPC.

**Figure 8.**
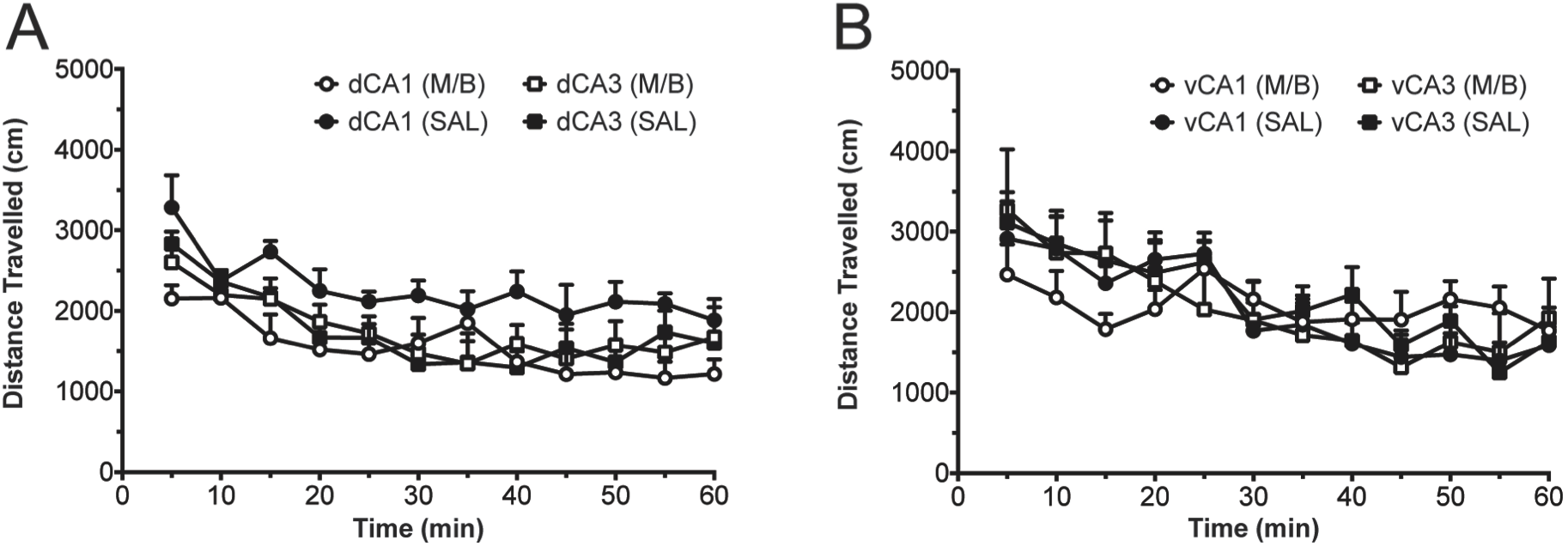
Mean (± SEM) distance travelled shown in 5 minute time bins during a test of general locomotor activity following administration of muscimol/baclofen (M/B) or saline (SAL) in the (A) dorsal CA1 and CA3 hippocampus, and in the (B) ventral CA1 and CA3 hippocampus. Mean and standard error values, as well as 95% confidence intervals are reported in Supplemental Table S5.

## DISCUSSION

To our knowledge, we have demonstrated, for the first time, that the vCA1 and vCA3 subregions of the rodent HPC make differential contributions to approach-avoidance conflict processing. Using a non-spatial, learned approach-avoidance paradigm, post-training GABAR agonist-mediated inactivation of vCA1 was found to increase avoidance of a cue associated with conflicting valence information whereas inactivation of the vCA3 led to potentiated approach behaviour in the face of motivational conflict. Notably, inactivation of dCA1 and dCA3 had no effect on conflict behaviour. Thus, in keeping with a large body of literature implicating a role for the ventral, but not dorsal, HPC in approach-avoidance conflict processing (Bannerman et al., 2004; 2012; Ito and Lee, 2016) our findings pertaining to the contrasting roles of CA1 and CA3 appear to be specific to the ventral portion of the HPC.

Previous insight into the role of ventral HPC subregions in approach-avoidance conflict processing has been predominantly limited to studies that have focused on the role of DG in anxiety, and which have yielded somewhat inconsistent results. Weeden et al. (2015) observed increased amount of time spent in the open arms of the elevated-plus maze as well as the centre of the open field test following selective lesions of the ventral DG (vDG), but not dorsal DG. In contrast, Kheirbek et al. (2013) reported that light-induced *activation* of the ventral DG granule cells led to a reduction in anxiety-related behaviour, using the same two ethological tests of anxiety, which could not be accounted for by a change in locomotor exploratory behaviour. Although it is currently unclear how differential manipulations of vDG (i.e. optogenetic activation vs. lesion) can lead to similar anxiolytic effects, these data collectively implicate a role for the vDG in regulating approach-avoidance behaviour under circumstances of innate conflict (e.g. preference for enclosed spaces vs. desire to explore in the elevated plus maze).

Our current findings add significantly to this recent work by revealing how other regions within the ventral HPC trisynaptic circuit contribute to approach-avoidance conflict processing, with the vCA3 and vCA1 appearing to play opposing roles. Not dissimilar to the aforementioned findings pertaining to vDG, inactivation of vCA3 was observed to increase the amount of time that rodents spent exploring the maze arm containing the conflict cue, demonstrating potentiated approach tendency in the face of learned motivational conflict. In contrast, vCA1 activation led to the opposite behavioral tendency (avoidance), with rodents spending significantly less time in the conflict cue arm and spending a higher proportion of time in the neutral cue arm. Given that previous findings from our laboratory have shown that ventral HPC lesions do not impair the acquisition of conditioned approach or avoidance behavior (Schumacher et al., 2016), the present results point to the vCA3 having a critical role in opposing/suppressing approach tendencies specifically in situations of learned approach-avoidance conflict, and the vCA1 in promoting approach behaviour under such conflict. This postulated role of the vCA3 is consistent with the findings of a recent optogenetic and chemogenetic study that observed suppression in feeding and anxiogenesis when the excitatory neurons in the vDG/vCA3 were chemogenetically activated, and conversely, facilitation of feeding when the vDG/vCA3 neurons were chemogenetically inactivated (Sweeney and Yang, 2015). Together with the plethora of evidence implicating the ventral HPC in suppressing approach responses in the face of a threat to energy homeostasis (Tracy et al., 2001; Davidson et al., 2009), the present findings help solidify the notion that the ventral HPC is fundamentally important in the regulation of innate and learned approach-avoidance decisions in states of environmental uncertainty and instability. Furthermore, the present study brings a novel extension of this view, in proposing that the ventral HPC has bidirectional control over approach-avoidance behaviours via region-specific, CA3-Aversus CA1-mediated mechanisms.

We speculate that such bidirectional control may be achieved through the vCA3 and vCA1 subfields operating as parts of independent circuits, as opposed to functioning in a serial fashion through a trisynaptic circuit (DG->CA3->CA1), in contrast to the traditional understanding of information flow through the HPC. In fact, the notion that each subregion does not necessarily depend on intrinsic circuitry for serial input is illustrated in studies in which pharmacological disruptions to the DG or CA3 are shown not to have any debilitating effect on place field activity in the CA1 (Mizumori et al., 1989; Brun et al., 2002). Furthermore, the differential pattern of extrinsic CA3 and CA1 connectivity provides the means by which CA3 and CA1 subregions can function independently of one another. While the CA3 is most known for its intrinsic excitatory associational (CA3-to-CA3) and commissural (CA3 - contralateral CA3 and CA1) connections that constitute an ‘auto-associational’ network that enables the rapid and efficient encoding and recall of information (Nakazawa et al., 2002; Van Strien et al., 2009), there is compelling neuroanatomical evidence to suggest that it has a robust extrinsic connectivity with the lateral septum (LS). The projections from the CA3 to LS are thought to occur in a topographical manner, with the dCA3 projecting to the dorsal aspects of the lateral septum, and the vCA3 projecting to more ventral parts of the lateral septum (Witter, 2007). The LS itself has been widely implicated in the regulation of anxiety, albeit the exact nature of its role remains undetermined due to the varied direction of effects that LS manipulations have produced. For instance, selective pharmacological inactivation of the LS has been shown to reduce anxiety in ethological tests of anxiety such as the elevated plus maze (EPM), and shock-probe burying test (Menard and Treit, 1996; Degroot et al., 2001). However, these findings are hard to reconcile with lesion studies that have reported ‘septal or sham rage’ - increased display of defensive behaviors to otherwise innocuous stimuli following lateral (and medial) septum lesions (Albert and Brayley, 1979; Blanchard et al., 1979), or studies which report reduced anxiety-like behavior when the LS is electrically stimulated (Yadin et al., 1993). Recent studies employing circuit-specific approaches have sought to further elucidate the role of the ventral HPC-lateral septal pathway in anxiety and feeding regulation, but have yielded inconsistent results. For instance, Parfitt et al., (2017) found that a chemogenetic activation of the LS-projecting ventral HPC cells led to reduction in anxiety, as tested in an array of ethological tests such as EPM and successive alley, while inactivation of the same neurons led to anxiogenic effects. In contrast, Sweeney and Yang (2015) found that optogenetic and chemogenetic activation of glutamatergic ventral HPC -> LS neurons suppressed food intake, while inactivation of lateral septal neurons blocked HPC-mediated suppression of feeding. Crucially, it should be noted that neither of these studies can confirm the exact locus of origin of the ventral HPC neurons projecting to the LS (e.g. CA3 vs. CA1), and the present findings highlight the importance of selectively targeting LS-projecting vCA1 and vCA3 neurons in future investigations.

In contrast to the CA3, the CA1 has a much wider extrinsic connectivity, projecting to a number of subcortical and cortical areas in addition to the LS (Van Groen and Wyss, 1990; Witter and Amaral, 2004). The LS-projecting CA1 neurons are also arranged topographically along the dorsal-ventral axis as with the projections originating in the CA3, although it is thought that the CA1 neurons terminate in more rostral areas of the LS compared to the CA3 neurons (Risold and Swanson, 1997; Naber and Witter, 1998). Thus, it is plausible that approach/avoidance behaviors are subserved by functionally separate, parallel ventral HPC-lateral septal loops. The CA1 also has extensive projections to the prelimbic/infralimbic cortex, amygdala and nucleus accumbens (NAc), and together with recent evidence from our laboratory demonstrating that transient GABAR_A&B_ receptor-mediated inactivation of the caudal NAc core induces the same effect on learned approach-avoidance decision making as the present vCA1 inactivation effect (potentiated conditioned avoidance) (Hamel et al., 2017), the CA1 and NAc core may be candidates structures for forming a functional pathway that facilitates approach behaviour in the face of environmental uncertainty. This possibility warrants further investigation.

One further significant advance that the present study makes is in moving beyond the domain of innate behaviour as assessed by ethological tests of anxiety, to examine HPC subregion contributions to approach-avoidance conflict that arise as a result of learned cue-valence associations. This is an important step since a state of approach-avoidance conflict can often arise in response to stimuli that have no innate value, and for which the associated valences are acquired over time. Notably, we also administered a classic ethological anxiety test, the light-dark box in the current study and found that the pattern of results in the test did not recapitulate the results obtained with the learned approach-avoidance conflict test. Inactivation of both the vCA 1 and vCA3 regions reduced anxiety, with a visual inspection of the graph depicting the vCA3 inactivation to have had a larger effect, with the rats spending more time in the more anxiogenic bright light box, as compared to the dark box. In contrast, the vCA1-inactivated rats appeared to spend equal time in the light and dark compartments. The inability to observe a direct correspondence in our vCA1 and vCA3 inactivation findings across the approach-avoidance conflict task and the dark-light box suggest that innate anxiety and learned approach-avoidance decision making are two dissociable psychological constructs that share some, and not all common neural substrates. In support of this, we have previously observed the manifestation of alterations in learned approach-avoidance conflict behaviour in the absence of concomitant changes in indices of innate anxiety following NAc core inactivation and repeated cocaine exposure (Nguyen et al., 2015; Hamel et al., 2017). We speculate that while approach-avoidance conflict processing may be a key component of anxiety, it is not the only contributing factor and that dysregulation of other decision-making and motivational processes are likely to contribute to the full spectrum of anxiety-related behaviour.

It is important to emphasize that the observed pattern of findings across HPC subregions and along the longitudinal axis cannot be accounted for by other factors including differences in cue acquisition (as discussed earlier), novelty detection or changes to locomotor activity. Firstly, we did not observe any significant changes in spontaneous activity in any of the ventral or dorsal HPC inactivation groups. We also failed to see any differences in the total number of entries made into the conflict and neutral arms between any groups during the conflict test, a measure that is typically sensitive to changes in baseline locomotor activity. Secondly, the fact that inactivation of *all* dorsal and vCA 1 and vCA3 subregions led to a marked impairment in novelty preference cannot fully explain the differential effect of manipulating the ventral and dorsal HPC CA3 and CA1 on learned approach-avoidance conflict behavior. Previous studies have implicated the dorsal HPC to be involved in spatial novelty processing (Lee et al., 2005; Wells et al., 2013), with one potential caveat that the successful novelty detection/preference requires the intact capacity to process spatial cues, which is also a function that the dorsal HPC is critical in (Moser et al., 1993; Bannerman et al., 1999; 2002). In the present task, we ensured that animals would be able to make use of both spatial (extra-maze) and non-spatial (intra-maze) cues to perform the novelty preference task, so we could assess the role of the dorsal and ventral HPC subregions in novelty processing *per se*, and to minimise the potential contribution of confounding factors (impaired spatial/cue processing). Very few studies have directly examined the role of the ventral HPC in novelty processing (Riaz et al., 2017), but the present findings suggest that the ventral HPC CA3 and CA1subregions are as important as the dorsal HPC in mediating novelty processing.

In conclusion, we have provided novel insight into the differential contributions of ventral HPC regions to learned approach-avoidance conflict processing. Specifically, ventral, but not dorsal, CA1 and CA3 appear to play opposing roles in the regulation of with the former facilitating approach and the latter avoidance when an animal is confronted with circumstances of high motivational conflict. Our findings have implications for our current understanding of the role of the HPC in motivational decision making and highlight the importance of considering differences not only along the longitudinal axis but also transverse axis of this structure. Furthermore, the observed contrasting effects of ventral CA1 and ventral CA3 inactivation upon approach-avoidance conflict behavior point to the existence of functionally distinct, extra-hippocampal neural circuits associated with individual HPC subfields, and thereby provide new insight into the functions and circuitry of the HPC beyond the much-studied unidirectional tri-synaptic hippocampal circuit.

## METHODS

### Subjects

Subjects were 80 male Long Evans rats (Charles Rivers Laboratories, QC, Canada) weighing between 350-400g prior to any procedures. Rats were housed in pairs with a constant room temperature of 21°C, under a 12 hour light/dark cycle. Water was provided *ad libitum* but food was restricted to maintain the rats at 85% of their free feeding body weight. All behavioral testing took place during the light cycle, in accordance with the ethical and legal requirements under Ontario's Animals for Research Act, the federal Canadian Council on Animal Care, and approval of the University of Toronto Scarborough Local Animal Care Committee.

### Surgery

All rats were surgically implanted with bilateral guide cannulae overlying the dorsal or ventral HPC CA3 or CA 1 regions, prior to the commencement of behavioral testing. Rats were assigned into 2 (ventral or dorsal HPC) x 2 (CA3 or CA1) x 2 (Drug vs. Saline) experimental groups according to the anatomical location of their implanted cannulae as well as the drug with which they would be injected with, that is, a muscimol (GABA_a_ receptor agonist) and baclofen (GABA_/b_ receptor agonist) cocktail (M/B) or saline (SAL): vCA1(M/B, n=12), vCA3(M/B, n=12), vCA1 (SAL, n=12), vCA3(SAL, n=12), dCA1(M/B, n=8), dCA3(M/B, n=8), dCA1(SAL, n=8), and dCA3(SAL, n=8). All operated rats were anesthetized with isoflurane gas and placed in a stereotaxic frame (Stoelting, IL, USA). A midline incision along the skull was made, and the fascia retracted by small skin clips to reveal the cranial landmarks lambda and bregma. Guide cannulae (23 gauge; Coopers Needle Works, UK) were then implanted bilaterally relative to bregma, targeting the dCA1 (AP -3.6mm; ML ±2.5mm; DV -1.8mm), dCA3 (AP -3.6mm; ML ±2mm; DV -2.4mm), vCA1 (AP -5.8mm; ML ±5.4mm; DV -6.5mm) and the vCA3 (AP -5.8mm; ML ±4.6mm; DV -5mm) in accordance with Paxinos and Watson (1998). Cannulae were anchored to the skull using dental cement (Lang Dental, IL, USA) and miniature stainless steel screws. Solid stainless steel dummy cannulae (30 gauge; Coopers Needle Works, UK) were inserted into the guide cannulae to ensure patency for the duration of the experiment. Rats were given 7 days to recover before any behavioral testing with water and food available *ad libitum*.

### Microinfusion Procedure

Muscimol and baclofen solutions were prepared separately at a concentration of 500ng/μl and combined in equal volumes to achieve a final concentration of 250ng/μl for each compound in accordance with previously reported dosages that induced behavioral alterations (Hamel et al., 2017; Riaz et al., 2017). The final infusion dose of the GABA_A/B_ receptor agonist cocktail was 75ng, delivered bilaterally at a volume of 0.3μl/side/minute, using an infusion pump (Harvard

Apparatus, Holliston, MA) mounted with a 5 μl Hamilton syringe. A recent finding from our laboratory (Hamel et al., 2017) revealed that a 0.3μl (75ng) infusion of Muscimol/Baclofen induced a discrete 0.3mm radial drug spread/inhibition in the target brain area (nucleus accumbens), as evidenced by a significant reduction in C-Fos activation in the drug-infused, as compared to saline-infused brains. Given that we have used the same dose and volume of muscimol and baclofen in the present study, we have high confidence in the fact that the spread of drug/active radius of inactivation remained well within the confines of targeted subfields (CA3 vs. CA1).

24 hours prior to the first drug infusion, all rats received a single infusion of 0.9% saline (SAL) bilaterally at 0.3μL/side to acclimatize rats to the infusion procedure, and to minimize the mechanical effects of subsequent drug infusions. During the infusion procedure, rats were lightly restrained, and the stainless-steel dummy cannulae were replaced with 30-gauge injectors (Plastics One, VA, USA) that extended 1mm beyond the guide cannulae. Injectors were connected to a syringe pump (WPI, FL, USA) that infused 0.3 μL of the drug cocktail or 0.9% saline, over 1 minute. Injectors were left in place for an additional minute to allow for diffusion of the drug/saline away from the injection site. Rats were returned to their home cage for 10-15 minutes before behavioral testing commenced.

### Behavioral Procedures

### Approach-Avoidance Conflict Task

#### Radial Arm Maze Apparatus

Behavioral testing for the approach-avoidance conflict task was performed in an automated six-arm radial maze as previously described (Med Associates, VT, USA) (Nguyen et al., 2015; Schumacher et al., 2016). Six identical enclosed arms (45.7 cm length X 16.5 cm height X 9.0 cm width) emanated from a hexagonal central hub, but only three out of the six arms were used throughout testing. Arms contained stainless steel grid floors connected to a foot shock generator and were enclosed by Plexiglas walls and removable lids covered in their entirety with red cellophane to limit the visibility and use of extra-maze cues. Automated stainless steel guillotine doors permitted access to the arms from the hub and vice-versa. The ends of each arm contained a port with a fluid receptacle connected to a syringe that allowed for the delivery of a 20% sucrose solution. A camera mounted above the apparatus was used to record behavioral testing. At the end of each session, the maze was cleaned with ethanol solution to eliminate odor traces and was rotated 60° clockwise to minimize conditioning to extraneous intra-maze cues.

#### Preconditioning Habituation

Rats were given three habituation sessions, as previously described (Schumacher et al., 2016). In each session, animals were placed in the central hub for one minute, followed by the opening of all two or three guillotine doors to allow the rats to freely explore the arms for a further five minutes. In the first habituation session, the rats were exposed to all three arms without any cue inserts. In the second habituation session, rats were exposed to three pairs of bar cues (45cm length x 4cm width x 0.5cm height, wood panels varying in color and texture) lining the full length of the sidewall of each of the arms. In this session, the exploration time of each cued arm was recorded to help determine the assignment of valence (appetitive, aversive, neutral) to each cue. Where there were innate preferences for one cue over the others, the most preferred cue was assigned the aversive valence, and the least preferred cue assigned the appetitive valence. During the third habituation session, rats were presented with two sets of cues in two arms, with one of the arms containing a pair of ‘to be assigned’ neutral cues, as determined from the second habituation session. The other arm contained a combinatorial cue comprised of one bar cue to be associated with appetitive valence and another bar cue to be assigned aversive valence. This session mimicked the conditions of the approach-avoidance conflict test (see section ‘Approach-Avoidance Conflict Test’) in order to eliminate the novelty of experiencing a combinatorial cue. Time spent exploring each cued arm was measured.

#### Non-spatial Mixed Valence Cue Conditioning

Cue conditioning sessions were conducted once per day over the course of nine consecutive days. In each conditioning session, rats were first placed in the central hub for 30 seconds followed by two minutes of confinement in each of the three cued arms, with the order of arm presentation counterbalanced across animals, and across sessions. In the arm containing the appetitive cue, rats received four randomly administered aliquots of 0.4ml of 20% sucrose solution, while in the arm with the aversive cue, rats were administered four mild foot shocks (0.5s, 0.25mA - 0.30mA) administered at a random inter-shock interval ranging from 15 - 25s. In the arm that contained the neutral cue, rats did not experience any reward or shock. Notably, previous work in our lab (Nguyen et al., 2015; Ito and Lee, 2016; Schumacher et al., 2016; Hamel et al., 2017) had established that these specific magnitudes of unconditioned stimuli (sucrose and foot shock) were required for the uniform, and balanced development of conditioned approach and avoidance behaviour. To ensure that outcomes were conditioned specifically to the bar cues (and not to any other available intra-maze or extra-maze cues), the placement of the bar cues was counterbalanced across rats, and changed across sessions, and the maze was rotated left or right by varying degrees (60°, 120°, or 180°) between each conditioning session. The entire maze was also covered with red cellophane film to block the visibility of extra-maze cues, while allowing video recording to take place via an infrared camera mounted on the ceiling.

#### Conditioned Cue Approach/Avoidance Test

Two conditioned cue approach/avoidance tests were conducted, one prior to conditioning session five and another prior to session nine, to demonstrate that the rats had learned the association between cues and their respective outcomes. The testing was identical to habituation session two, with the rats allowed to explore the appetitive, aversive, and neutral cued arms in extinction (without any outcomes) for 5 minutes. The time spent exploring each arm was recorded for each test. Successful acquisition of the cue contingencies was determined as rats spending more time exploring the appetitive cue (conditioned approach) and less time exploring the aversive cue (conditioned avoidance), relative to the neutral cue.

#### Approach-Avoidance Conflict Test

Prior to the approach-avoidance conflict test, rats that demonstrated successful cue acquisition underwent drug or saline infusion into the target hippocampal area (see section ‘Microinfusion Procedure’). During the conflict test (in which 2 maze arms were used), rats were first placed in the central hub for one minute, after which a state of approach-avoidance conflict was induced by presenting the appetitive and aversive cue concurrently in one arm, and presenting the neutral cue in another arm. During this test, two measures were recorded: 1) the total time spent in the conflict arm and neutral arm; and 2) the number of full bodies entries made into each of the two arms.

### Light Dark Box

The light dark box test, used as a measure for innate anxiety, was conducted immediately after the rat finished the approach-avoidance conflict test while the drug effect was still likely to be present. The apparatus consisted of two conjoined compartments (Plexiglas; 60cm length x 30cm width x 25cm height), one with transparent walls (light box), and another with opaque black walls (dark box). An opaque black divider separated the compartments with an opening (12cm width x 12cm height) at the center of its base that allowed access between the compartments. The light box was illuminated by a lamp (11 watts) hanging 15cm above the ceiling of the light compartment. Both compartments were sealed using a wire mesh, and the dark box was covered by an opaque black Plexiglas sheet to prevent light entry. During the test, rats were placed in the middle of the light box and given 10 minutes to freely explore the apparatus. Time spent in each box was recorded. Furthermore, the light-dark ratio, calculated by the time spent in the light box relative to the total time in both boxes, was used to show differences between the inactivated CA1 and CA3 groups compared to their control groups.

### Novelty Detection

The same 6-arm radial maze from the approach-avoidance conflict task was used to test novelty detection in rats. Three of the six arms that were not used in the approach-avoidance conflict test were used and decorated with distinct visual cues lining outside the arm walls, and the lids were left open for visual access to extra-maze cues. Rats were therefore able to use both intra-and extra-maze cues to detect novelty. Prior to the novelty detection test, rats underwent the drug/saline microinfusion procedure (see section ‘Microinfusion Procedure’). The test consisted of two phases: a habituation and a test phase. During habituation, rats were placed at the end of one arm and presented with an additional arm. Rats were permitted to explore both (familiar) arms for 10 minutes, and the time spent exploring each arm was recorded. If the rats showed similar exploration pattern for both arms, they were tested in the second and final phase. During the test phase, rats were given access to a third “novel” arm and to the two familiar arms for 5 minutes. Time spent exploring each arm was recorded, and an average for the time spent exploring the two familiar arms was calculated for comparison with the novel arm.

### Locomotor Activity

The locomotor activity test was conducted following completion of the novelty detection test while rats were still under the influence of the drug. Rats were placed individually in activity chambers (44cm length x 24cm width x 20cm height) lined with standard bedding and sealed with stainless steel chamber lids. Total distance travelled (in cm) for one hour, divided in 12 five-minute bins, was measured using an overhead camera and EthoVision tracking software (Noldus Information Technology, ON, Canada).

### Histology

All rats were deeply anaesthetized with an overdose of pentobarbital (200 mg/kg intraperitoneal; Bimeda-MTC, ON, Canada) and transcardially perfused with 0.9% saline and 4% paraformaldehyde. Brains were removed and stored overnight in 4% paraformaldehyde, followed by 30% sucrose for an additional 48 hours. Brain tissue was then frozen and sliced (50μm) using a freezing microtome, mounted onto slides and stained with cresyl violet, to be viewed under the microscope for verification of correct cannula and injector tip placement. Based on the Paxinos and Watson brain atlas (1998), rats with misplaced cannulae and injector tips that extended beyond the boundaries of each hippocampal subfield were excluded from the study (see section ‘Histological Verifications’ for details).

### Statistical Analysis

Data were analyzed using the SPSS statistical package version 21.0 (IBM, ON, Canada). Analysis of variance (ANOVA) was applied to all experimental data. The factors “region” and “drug” were set as the between-subjects factors for all behavioral tasks, while the factor “arm” served as the within-subjects factor for the habituation and conflict tests of the approach-avoidance conflict task, as well as the light dark box (referred to as ‘compartment’), and novelty detection. The locomotor activity task was analyzed with the factor “bin” as the within-subjects factor, while both acquisition tests of the approach-avoidance conflict task were analyzed with the factors “arm” and test” as within-subjects factors. Furthermore, all significant main within-subjects effects, three- or four-way interactions were explored further using paired samples t-tests, simple effect analyses, and/or post-hoc comparisons with a Bonferroni correction.

## COMPETING INTERESTS

The authors have no competing interests to declare.

